# Detection of structural mosaicism from targeted and whole-genome sequencing data

**DOI:** 10.1101/062620

**Authors:** Daniel A. King, Alejandro Sifrim, Tomas W. Fitzgerald, Raheleh Rahbari, Emma Hobson, Tessa Homfray, Sahar Mansour, Sarju G. Mehta, Mohammed Shehla, Susan E. Tomkins, Pradeep C. Vasudevan, Matthew E. Hurles, The Deciphering Developmental Disorders Study

## Abstract

Structural mosaic abnormalities are large post-zygotic mutations present in a subset of cells and have been implicated in developmental disorders and cancer. Such mutations have been conventionally assessed in clinical diagnostics using cytogenetic or microarray testing. Modern disease studies rely heavily on exome sequencing, yet an adequate method for the detection of structural mosaicism using targeted sequencing data is lacking. Here, we present a method, called MrMosaic, to detect structural mosaic abnormalities using deviations in allele fraction and read coverage from next generation sequencing data. Whole-exome sequencing (WES) and whole-genome sequencing (WGS) simulations were used to calculate detection performance across a range of mosaic event sizes, types, clonalities, and sequencing depths. The tool was applied to 4,911 patients with undiagnosed developmental disorders, and 11 events in 9 patients were detected. In 8 of 11 cases, mosaicism was observed in saliva but not blood, suggesting that assaying blood alone would miss a large fraction, possibly more than 50%, of mosaic diagnostic chromosomal rearrangements.

## INTRODUCTION

Genetic mutations that arise post-zygotically lead to genetic heterogeneity in an organism, a phenomenon called mosaicism. The detection of mosaic mutations that are small (single-base or indel) is still a great technical challenge but multi-megabase (“structural“) mosaic rearrangements are now routinely detected using cytogenetics and microarray technology (Miller et al. 2010; Biesecker and Spinner 2013). Recent single nucleotide polymorphism (SNP) microarray-based studies have demonstrated that mosaic structural abnormalities are implicated in developmental disorders (Conlin et al. 2010; King et al. 2015), increase in incidence with age (Forsberg et al. 2012), and predispose to hematological malignancies in adults (Jacobs et al. 2012; Laurie et al. 2012).

Modern SNP microarray technology is well suited for detecting mosaicism because probe density is high (often above 1 million sites per genome) and probes generate allele ratio data with high signal to noise ratio. SNP microarray platforms assess two metrics useful for mosaicism detection: 1) b allele frequency (BAF): the fraction of the alleles at a locus representing the less-common allele and 2) log R ratio (LRR): a measure of copy-number, based on the log ratio of signal intensity compared to a reference. These metrics are affected differently depending on the nature of the structural abnormality: whereas copy-neutral (loss of heterozygosity; LOH) mosaicism results in a deviation of BAF alone, copy-number (gain or loss) mosaicism additionally alters the LRR. Absolute deviation from genotype-expected BAF (e.g. 0.5 for AB genotype), called B-deviation (B_dev_), occurs in mosaic regions when the locus has a mixture of genotypes from wild type and mosaic tissue. Several software tools (Partek^®^ Genomics Suite, Illumina^®^ cnvPartition, BAFsegmentation (Staaf et al. 2008), and Mosaic Alteration Detection (MAD) (González et al. 2011)) harness this deviation as a mosaic signal. MAD is open source and has been recently used in several large SNP-based mosaicism projects (Forsberg et al. 2012; Jacobs et al. 2012; Forsberg et al. 2014); it identifies mosaic segments using aberrations in B_dev_ and then labels aberrant segments as copy-loss, copy-gain, or copy-neutral events based on the alteration of the LRR from baseline, a deviation referred to here as copy-deviation, or C_dev_.

Developmental disorders (DD) are often caused by rare, small (SNV and indel) mutations, genetic variation which is not easily captured using microarray King et al. 2014). Therefore, to achieve a more comprehensive assessment of pathogenic mutations, rare disease studies rely heavily on targeted sequencing of the protein-coding regions (‘exons’) of the genome, an approach called whole-exome sequencing (WES) (Koboldt et al. 2013). Indeed, sequencing of the whole genome (WGS) offers several advantages compared to WES, including greater breadth of the genome and more consistent coverage of exons (Meynert et al. 2014). However, WGS is not currently as widely used as WES for rare disease studies due to higher costs, so this work focuses primarily on exome-sequencing data.

In addition to small-scale variation, forms of large-scale ‘structural variation’, including copy-number (Lee et al. 2007) and copy-neutral variation (uniparental disomy (UPD)) (Yamazawa et al. 2010), are also important causes of DD. CNV burden analysis of nearly 16,000 children with DD (Cooper et al. 2011) demonstrated that nearly all CNVs greater than 2 Mb are likely pathogenic (odds ratios for CNVs of 1.5 Mb and 3 Mb were 20 and 50, respectively), and that deletion events are more often penetrant than duplication events. UPD events are only present in about 1 in 3,500 healthy individuals (Robinson 2000), but are enriched in children with DD (King et al. 2014), and may result in highly penetrant imprinting disorders, recessive diseases, or may be associated with chromosomal mosaicism (Eggermann et al. 2015). Low-clonality mosaicism is difficult to observe in karyotyping, as inspection of at least 10 cells is required to exclude 26% mosaicism with 95% confidence (Hook 1977), and is also difficult to observe in microarray analysis, as the detection sensitivity of mosaic duplications by SNP microarray with about 1 million probes for events of at least 2 Mb in size is limited to events of at least 20% clonality (González et al. 2011). The median average clonality in recent SNP-based studies of DD for mosaic aneuploidy was 40% (Conlin et al. 2010), and for mosaic structural variation (2 Mb and greater), was 44% (King et al. 2015). Among children investigated with clinical diagnostic testing, the frequency of autosomal mosaic *copy-neutral* events was 0.24% (12 in 5,000) (Bruno et al. 2011) and the frequency of autosomal mosaic *copy-number* events was 0.35% 36 in 10,362) Pham et al. 2014). Combining these frequencies yields a combined frequency among cases of 0.59% of mosaic structural variation.

The detection of large-scale mutations from WES data is challenging because input data are derived using sparse sampling of the genome, as targeted regions typically cover only about 2% of the genome (Meynert et al. 2014), and sequence read depth at exons is biased by enrichment efficiency and other factors (Plagnol et al. 2012). Despite these limitations, exome-based software tools have been successfully engineered to detect large-scale *constitutive* mutations, including copy-number variation (Magi et al. 2013; Sathirapongsasuti et al. 2011; Krumm et al. 2012; Backenroth et al. 2014; Fromer et al. 2012) and copy-neutral variation (bcftools roh (Narasimhan et al. 2016) and UPDio (King et al. 2014)). These tools are relatively insensitive to *mosaic* abnormalities, however, because they typically rely on single metrics, such as copy-number change (rather than copy-number *and* allele-fraction), or on genotype, which is not well assessed in mosaic state. Specialized methods have been developed for the analysis of cancer exomes where tumor and normal tissue can be isolated (Lonigro et al. 2011; Amarasinghe et al. 2014) or, in the context of a parent-fetus trio, for fetal DNA in maternal plasma (Rampášek et al. 2014). However, a method to detect copy-number and copy-neutral mosaicism from an individual’s exome (or genome) is lacking, but if available, could further extend the capacity of sequence-based analyses.

We developed MrMosaic, a method that detects structural mosaicism using joint analysis of B_dev_ and C_dev_ in targeted or whole-genome sequencing data (Figure 1). We used simulations to demonstrate the superior performance of MrMosaic compared to the MAD algorithm. We also applied MrMosaic to analyze WES data from 4,911 children with developmental disorders and identified 11 structural mosaic events in 9 individuals, 6 of whom exhibited tissue-specific mosaicism.

**Figure 1:**
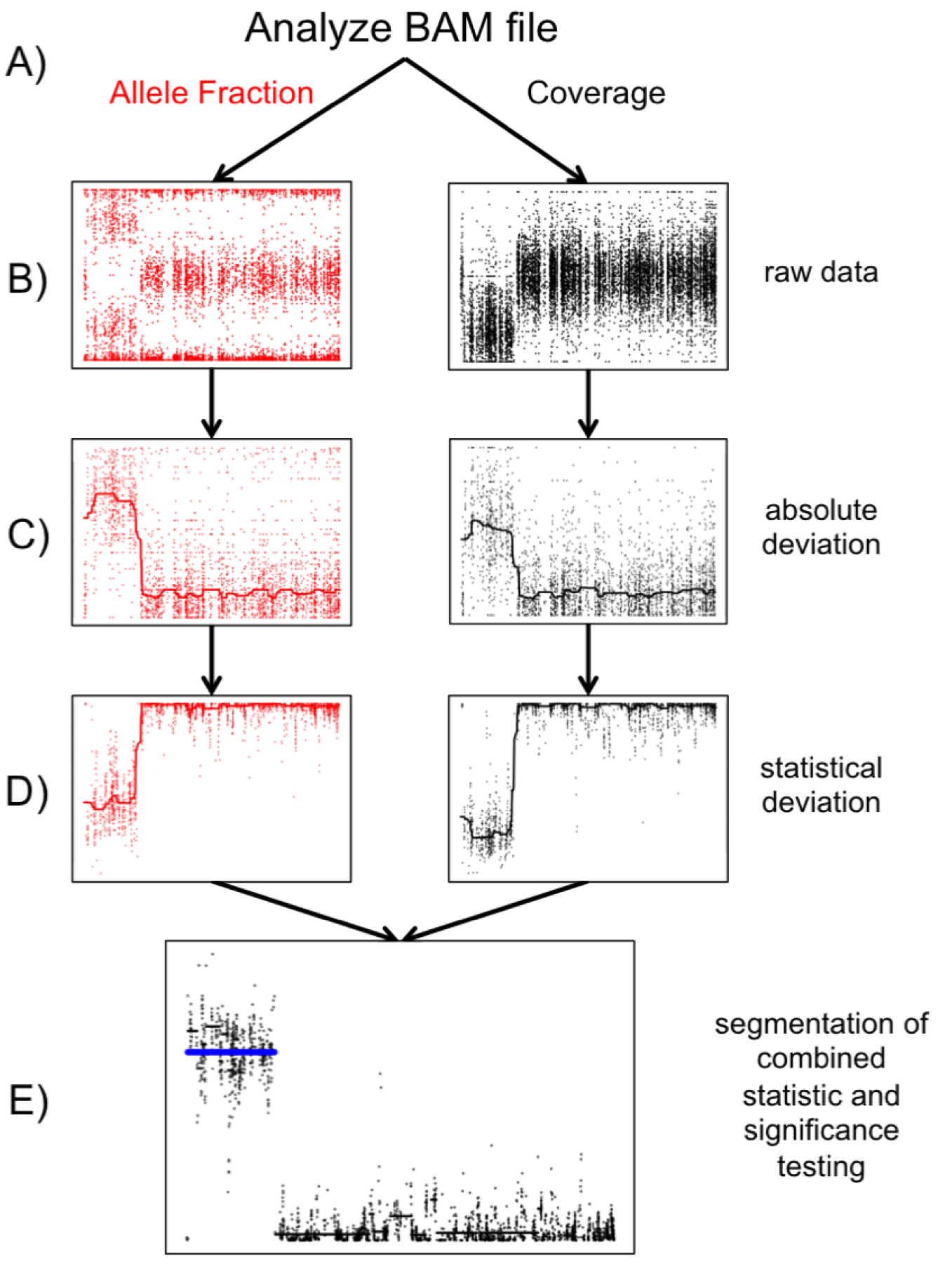
Detecting structural mosaicism using MrMosaic: A) Exome data are stored in a BAM file from which allele fraction (left column) and coverage (right column) are measured at polymorphic positions within or near target regions. A simulated mosaic deletion is depicted. B) The raw data, consisting of BAFs and normalized coverage are plotted for a simulated mosaic deletion. C) Absolute deviation of BAF (B_dev_) and normalized coverage (C_dev_) at heterozygous sites are analyzed. D) Mann Whitney U Tests are performed separately for B_dev_ and C_dev_, comparing the signal detected in sliding windows in this chromosome, compared with a randomly selected chromosome for background. E) The test statistics are depicted on the log scale. The p values of the Mann Whitney U Tests are combined and segmented (black lines). Segments passing the Mscore significance threshold are plotted in blue.

## RESULTS

We developed a new computational method, MrMosaic, to detect structural mosaic abnormalities from high-throughput sequence data (Methods). In summary, this method identifies chromosomal segments with elevated deviations in allelic proportion and copy number, relative to randomly selected sites on other chromosomes from the same data (Figure 1). Initially, measures of deviation of allelic proportion (B_dev_) and copy number (C_dev_) are computed from the WES/WGS data at well-covered known polymorphic SNVs. Whereas B_dev_ is only assessed at heterozygous sites, C_dev_ integrates information from flanking non-heterozygous sites to reduce noise. The statistical significance of the observed B_dev_ and C_dev_ are assessed separately, using non-parametric testing, and the resultant p values are subsequently combined and then segmented using the GADA algorithm (Pique-Regi et al. 2008). We devised a confidence score, the Mscore, to curate putative detections of mosaic segments, by integrating metrics that discriminate between true positive and false positive mosaic detections Methods).

## Simulations

We performed simulations (Methods) to explore the performance of MrMosaic for three different classes of structural mosaicism: gains, losses and LOH, in several contexts. The variation in performance across mosaicism of different *sizes, clonalities* and sequencing *coverage* is summarised in Figure 2, for both WES and WGS data.

**Figure 2:**
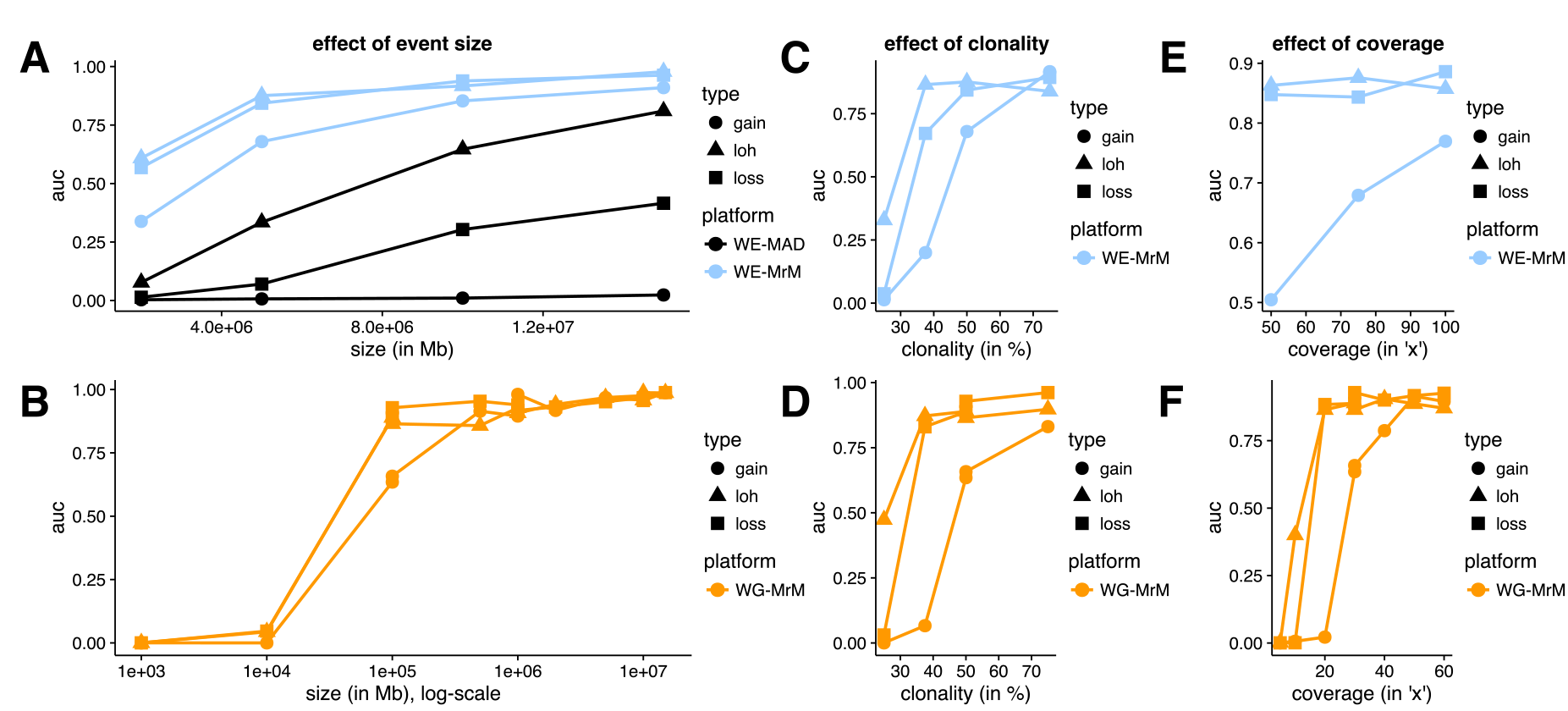
Simulation performance summarised by AUC: We measured the average precision (area under the precision recall curve) for MrMosaic implemented on whole-genome (WG) simulations (panels A,C,E), and MrMosaic & MAD implemented on whole-exome (WE) simulations (panels B,D,F). The depth, size, and coverage measured for WGS and WES simulations were selected to accentuate informative differences in performance. AUC across size: Simulated events of 50% clonality were studied for WGS (A) and WES (B) simulations. Whereas for WES simulations, simulated exome depth was 75x depth, for WGS simulations it was 30x depth. MrMosaic on whole-genome data (WG-MrM) outperforms MrMosaic on exome data (WE-MrM), which outperforms MAD on exome data (WE-MAD). AUC across clonality: Whereas for WES (C) simulations the simulated size and coverage was 5 Mb & 75x, for WGS (D) simulations it was 100 kb & 30x. AUC across average coverage: Simulated events of 50% were studied for both WES (E) and WGS (F) simulations. Whereas for WES simulations, simulated event size was 5 Mb, for WGS simulations it was 100 kb.

Across all measured categories, mosaic duplications were more difficult to identify than deletion or LOH events, especially at lower (25%) clonality (Supplementary Figure 1). We suspected that the most likely explanation for this lower sensitivity is that duplications result in the smallest deviation of B_dev_, compared with deletion and LOH events (Supplementary Figure 2) and that the C_dev_ signal is masked by sampling noise at low clonality. To further explore the effect of including C_dev_ in addition to B_dev_, we investigated the performance of MrMosaic using B_dev_ alone compared with joint analysis of B_dev_ and C_dev_. This analysis showed substantially improved detection of copy-number events above lower clonality, while only a marginally decreased performance of LOH detection (Supplementary Figure 3), consistent with the intuition that C_dev_ yields a valuable net signal when clonality is above the C_dev_ noise floor.

Simulation performance increased with larger event *size* (Figure 2A). WES simulation analysis demonstrated high area under the precision-recall curve (AUC) for all events at least 10 Mb in size and at least 50% in clonality; and, for deletion and loss of heterozygosity (LOH) events at least 5 Mb in size. MrMosaic performed favourably compared to MAD in all measured categories. Results for WGS simulations demonstrated an AUC of about 0.9 for 100 kb LOH and loss events, and greater than 0.95 for all megabase-size events. Larger events were assayed by more positions, and whole-genome simulations interrogated nearly 50-fold more sites than exome data Supplementary Table 1).

Detection performance in simulations increased between 25% and 75% *clonality* (Figure 2B). The WES and WGS clonality performance results were measured at 5 Mb and 100 kb sizes, respectively, as events at these sizes were most sensitive to changes in clonality (Supplementary Figs. 4 and 5). Previous studies of children with DD have reported a median mosaicism of approximately 40% mosaicism and detection performance is strong for detecting mosaicism at this clonality at the studied sizes. As clonality increases, the mosaicism is present in a greater proportion of cells, resulting in a greater signal of detection.

Simulation performance increases with respect to sequencing *coverage* (Figure 2C). The WES and WGS performance with respect to sequencing coverage were assessed for events of 50% clonality, using 5 Mb events for the WES simulations, and 100 kb events for the WGS simulations. WES simulations demonstrated a marginal improvement of detection performance at higher coverage, which was notable for mid-clonality gains (Supplementary Figure 4). Previous work has suggested that 75x average coverage in WES data is sufficient for constitutive copy-number analysis^33^ and these coverage simulations demonstrated that this exome coverage is also sufficient for the detection of mosaic structural abnormalities. In the WGS results, AUC rose dramatically between 15x and 20x for LOH and loss events and between 25x and 30x for gains. AUC was above about 0.9 for LOH and loss events at 30x depth, a standard sequencing depth used in WGS disease studies. Nearly all structural mosaic events of 100 kb and 50% clonality were detected Supplementary Figure 5) and average coverage of 20x was sufficient to detect nearly all 50% clonality deletion and LOH events at 100 kb, while detection performance of gains improved at 30x and 40x (Supplementary Figure 6). This improved performance as coverage increases results primarily from sampling variance (‘noise’) decreasing (correlation r = −0.95; Supplementary Figure 7), with an additional minor contribution from more sites more signals) passing the minimal depth threshold for consideration Supplementary Table 1).

## Detections in 4911 case exomes

We generated WES data for 4,911 children with undiagnosed developmental disorders. DNA was collected from either blood (n=1652), saliva (n=3246) or both (n=13), and sequenced to a median average coverage of 90X. Analysis for structural mosaicism identified 11 mosaic abnormalities among 9 individuals, a frequency of 0.18%. The detections consisted of five losses (median size: 13 Mb, median clonality: 46%), four gains (median size: 25 Mb, median clonality: 55%), and two LOHs (median size: 50 Mb, median clonality: 26%) (Figure 3, Table 1, Supplementary Figs. 8-18).

**Figure 3:**
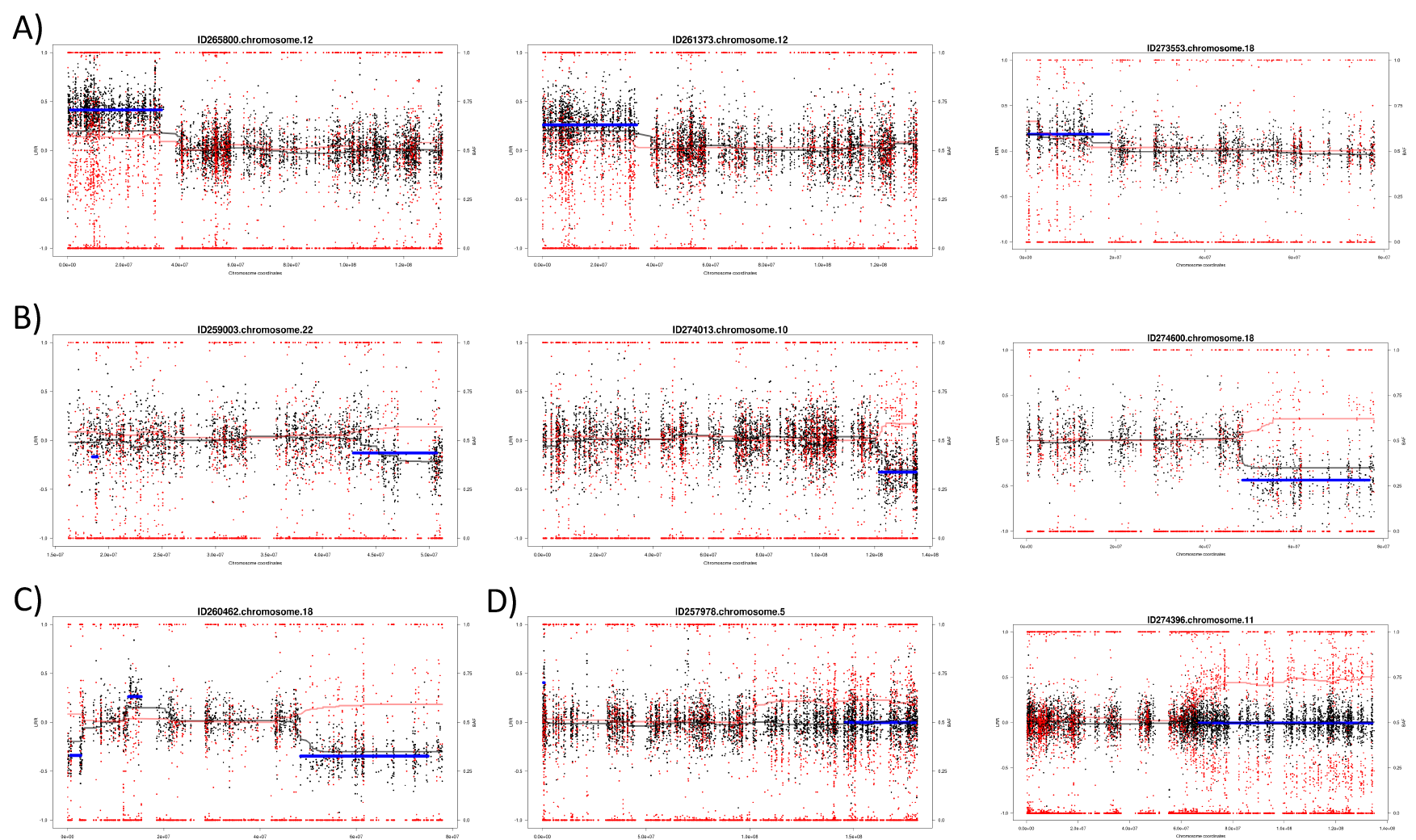
Structural mosaicism detected from exome data: Structural Mosaicism Detected by MrMosaic in the DDD study. Black and red dots represent copy-number and allele fraction, respectively. Cdev and Bdev are plotted in black and red trend lines. The blue line represents statistically significant segmented detections passing a threshold. Different classes of events are found: A) Mosaic gains, B) mosaic losses, C) mixed copy-number, and D) loss-of-heterozygosity events.

Previous analysis of a subset (1,226 of 4,911) of these samples by SNP microarray identified 10 events (King et al. 2015), while exome analysis yielded 8 events. Of the two events not detected by exome but detected by SNP microarray, one of the missed events was a 4 Mb duplication below 25% clonality. The other missed event was an LOH event with low sequencing depth (33x, one of the lowest of our study – Supplementary Figure 19); low depth results in higher sampling variance and lower statistical significance of deviations in allelic proportion and copy number (Supplementary Figure 7). Given the high clonality (about 75%) of this event, it may have been detected using constitutive (genotype-based) UPD analysis (although, as paternal data were not available for this sample, it was not analysed by our trio-based UPD detection pipeline (King et al. 2014)).

**Table 1:** Detections by exome and validation by SNP microarray: The 11 mosaic abnormalities detected in the 9 samples with exome data were validated using SNP microarray chips. All exome detections were validated in at least one tissue. In the majority of cases (8 of 11), the mutation was detected in only one of two assayed tissues, and in all such cases, the mutation was detected in saliva but not in blood. Clonality was calculated from Bdev using Equation 2 (see Supplementary Table 5) and ranged from 17% to 68%. This calculation is based on the assumption that the mosaic event is an alteration of a single allele. However, this calculated clonality is an overestimate for one of the events which was found (by previous FISH analysis^4^) to be a mosaic tetrasomy, and two others were are suspected to also be rearrangements of multiple alleles (another gain of chromosome 12p and one gain of chromosome 18p, thought to reflect mosaic tetrasomy 18).

| Exome Detections |  |  |  |  |  |  |  |  | SNP Validation |  |
| --- | --- | --- | --- | --- | --- | --- | --- | --- | --- | --- |
| DecipherID | chr | type | start<br>(GRCh37) | end<br>(GRCh37) | bdev | l2r | tissue | clonality | clonality<br>saliva | clonality<br>blood |
| 265800 | 12 | gain | 988,894 | 33,535,510 | 0.201 | 0.140 | saliva | 1.34 | 0.68<sup>@</sup> | absent |
| 261373 | 12 | gain | 283,642 | 33,535,289 | 0.131 | 0.262 | saliva | 0.72 | 0.45<sup>@</sup> | absent |
| 273553 | 18 | gain | 670,541 | 18,534,702 | 0.186 | 0.185 | saliva | 1.18 | 0.6<sup>@</sup> | absent |
| 259003 | 22 | loss | 42,912,136 | 50,717,129 | 0.131 | -0.129 | blood | 0.42 | 0.54 | 0.34 |
| 274013 | 10 | loss | 121,717,932 | 134,916,366 | 0.159 | -0.324 | saliva | 0.48 | 0.44 | absent |
| 274600 | 18 | loss | 48,458,662 | 76,870,586 | 0.190 | -0.434 | saliva | 0.55 | 0.49 | absent |
| 260462 | 18 | loss | 662,103 | 2,740,714 | 0.171 | -0.339 | saliva | 0.51 | 0.46 | absent |
| 260462* | 18 | gain | 12,702,610 | 15,323,214 | 0.118 | 0.263 | saliva | 0.41 | 0.5 | absent |
| 260462 | 18 | loss | 48,466,843 | 74,962,645 | 0.153 | -0.345 | saliva | 0.47 | 0.45 | absent |
| 257978 | 5 | LOH | 146,077,526 | 179,731,635 | 0.167 | -0.002 | blood | 0.33 | 0.24 | 0.26 |
| 274396 | 11 | LOH | 66,834,252 | 134,126,612 | 0.255 | -0.0047 | saliva | 0.51 | 0.28 | 0.17 |

<sup>@</sup>adjusted tetrasomy clonality.
\*located in peri-centromeric region and detected during *post hoc* analysis.

Validation of the 11 mosaic abnormalities using SNP microarrays on DNA derived from both blood and saliva successfully detected all abnormalities in at least one tissue (Table 1). Notably, six of the seven mosaic copy-number mutations detected by MrMosaic in exome data had been undetected by both clinical and high-resolution aCGH investigation of the same tissue, despite most events being at least 5 Mb in size and exhibiting 50% clonality (Supplementary Table 2). Examination of the raw aCGH data in one case (Supplementary Figure 17) showed that only small fragments of one of the events were detected but these called segments were individually much smaller than the actual event.

Detection of the mosaic events was largely dependent on the assayed tissue. Out of the 11 mosaic events, 3 were detected in blood and in saliva samples while the remaining eight were only observed in saliva (Table 1, Supplementary Figures 8-18). There were 2 abnormalities detected from 1,652 blood samples and 9 detected from 3,246 saliva samples, a non-significant proportional difference (p > 0.05, Fisher’s exact test). One of the mosaic events detected in both blood and saliva was an LOH-type event, remarkable for having a gradient of increasing clonality toward the telomere Supplementary Figs. 16 and 19). This gradient of increasing clonality along the chromosome is compatible with LOH-mediated mosaic reversion, characterised by distinct cell populations carrying partially overlapping independent LOH events, as reported recently (Choate et al. 2015). Nevertheless, despite generation and analysis of high-depth (~400x) WES data for this sample, and the identification of several strong candidate genes, including *CEP57* (the cause of mosaic aneuploidy syndrome (Snape et al. 2011)) in the reversion-localised region, no plausibly pathogenic rare below 1% minor allele frequency) coding sequence variants were identified Supplementary Table 4).

We assessed the pathogenicity of the events detected in these nine children based on overlap with known genomic disorders and diseases of imprinting Supplementary Material). The mosaic events identified in seven of nine children were considered definitely pathogenic on the basis of being multi-megabase CNVs that overlap known genomic-disorder regions. The reversion mosaic event was considered indicative of a likely pathogenic mutation as the presence of multiple overlapping mosaic clones suggests strong and on-going negative selection against a deleterious allele. One LOH event was of uncertain pathogenicity as no rare loss-of-function or functional variants were detected (Supplementary Table 4).

## Empirical evaluation of detection of mosaicism from WGS data

We selected one sample with three mosaic abnormalities detected on a single chromosome to demonstrate MrMosaic performance on whole-genome sequence data and to investigate the structure of the mosaic rearrangement. MrMosaic easily detected these multimegabase mosaic events, found with Mscores of 36, 117, and 32. The presence of three mosaic events of similar clonality on the same chromosome is suggestive of a complex chromosomal rearrangement. Analysis of the WGS read pair data using Breakdancer (Chen et al. 2009) identified read-pairs mapping across the centromere and evidence of a breakpoint spanning from the q-arm deletion to the centromere. Ring chromosomes are associated with bi-terminal deletions (Guilherme et al. 2011) and inverted duplications (Knijnenburg et al. 2007) and we suspected that the underlying abnormality in this child is a ring chromosome, although we were unable to access the cellular material required to generate the cytogenetic data to prove this hypothesis (Supplementary Figure 21).

## DISCUSSION

Structural mosaic abnormalities are multi-megabase, post-zygotic mutations that have previously been associated with developmental disorders (Conlin et al. 2010; King et al. 2015). This work introduces a novel method to detect these mutations from next generation sequencing data.

In an extensive simulation study we show adequate power to detect abnormalities in WES and WGS data across a large, clinically relevant range of size and clonality in different types of mosaic structural variation. We also compare our method to the popular array-based mosaic detection method, MAD, and show a substantial boost in performance, which derives primarily from the joint analysis of allelic proportion and copy-number deviations. Simulation results suggested that exome sequencing data can be used to identify many of the known clinical mosaic duplications involving chromosome-arm events, such as 12p and 18p mosaic tetrasomy as MrMosaic easily detected events of this size.

We used MrMosaic to uncover pathogenic structural mosaicism in a large exome study of children with undiagnosed developmental disorders. Applying our method to the exome data of 4,911 enrolled children, we identified nine individuals with structural mosaicism; the majority of these mutations were considered pathogenic. In this WES-based analysis we recovered 8 of the 10 abnormalities previously detected in a subset of 1,226 samples previously analysed with SNP genotyping chip data suggesting that exome-analysis alone is sensitive to detecting large-scale mosaicism. One of the missed abnormalities was likely undetected because the exome data were of low depth, which increases the variance of measured B_dev_ and C_dev_. Most of the detected mosaic copy number abnormalities had escaped detection by previous aCGH analysis. This demonstrates that detection of mosaic events requires assay of tissue containing the abnormality and tailored methods with sufficient sensitivity for mosaicism.

The overall frequency of mosaicism detected in this study, 0.18%, is lower and significantly different (p < 10^−4^, binomial test) from the 0.59% structural mosaicism frequency estimated from previous studies. One likely explanation for the discrepancy in these frequencies is ascertainment bias, as some classes of structural mosaicism (e.g. mosaic trisomies) are likely to have been diagnosed by prior diagnostic testing (e.g. karyotype or microarray) and not enrolled into the DDD study. Another component of this discordance may be due to decreased sensitivity, as mosaicism smaller than 2 Mb is challenging to detect by exome and these small events account for ~25% (9/36) of mosaic copy number events described previously (Pham et al. 2014).

In one sample we observed a gradient of mosaicism, a phenomenon likely associated with mosaic reversion of a *de novo* mutation inducing genome instability. Analysis of the mosaic LOH region with high-depth exome data did not identify a strong candidate coding variant and a further WGS-based search for candidate pathogenic *de novo* mutations is on-going. Whole genome sequencing data were generated for one individual with three mosaic abnormalities on the same chromosome. Analysis of these data recapitulated the mosaic events and analysis of read pair analysis identified a pericentromeric inversion and supported the hypothesis of an underlying complex chromosomal rearrangement, likely a ring chromosome.

As expected, whole genome analysis had superior performance compared to exome analysis, which was likely due to a combination of advantages of whole-genome data, including higher density of assayed sites by nearly 50 fold) and more consistent coverage across sites, compared to exome coverage, which is subject to exome bait hybridisation biases. Compared to whole genome data, the exome data had higher average coverage (75x to 25x) for sites within targeted regions compared to the whole genome data and while simulation results showed increasing performance with higher depth sequence data, this effect was outweighed by the greater density of sites in whole genome data.

Although the general performance of the method is adequate in many clinically-relevant cases, some classes of event prove more difficult to detect. For example, low clonality mosaic gains generate the smallest deviation in B_dev_ and C_dev_ compared to other types of events, explaining their comparatively poor detection sensitivity in simulations, and the failure to detect one mosaic duplication found using SNP data but not in exome data. More lenient detection thresholds may be preferred to increase detection sensitivity if clinical suspicion of mosaic duplication exists. Increasing the clonality of mosaicism by the biopsy of affected tissue, as is performed when pigmentary mosaicism provides evidence of underlying mosaicism Woods et al. 1994), should also theoretically improve detection. Given the size and clonality of the two missed events and the simulation results from whole genome sequencing, both events would likely have been detected had they been analysed using higher depth WES or WGS, which are likely to become more common in the future.

The majority of the mosaic events we observed in saliva-derived DNA were not observed in blood. The samples with these abnormalities were recruited into our study because they remained undiagnosed after assessment by clinical laboratories of blood-derived DNA failed to detect the mosaic abnormalities we detected in saliva. DNA derived from saliva has a mixed origin, mainly lymphocytes (derived from mesoderm) and epithelium (derived from epiderm) (Endler et al. 1999); therefore the events detected in saliva, but not blood, are believed to reflect epithelial mosaicism. There are two possible explanations for the disparity in tissue distribution we observed: first, that the epithelium-derived mutational events occurred late, i.e. after the differentiation of lymphocytes and epithelial cells, or second, that these events occurred early, i.e. prior to the split between lymphocytes and epithelial cells with subsequent removal from blood cell lineages by purifying selection. Several lines of evidence suggest the second explanation is more likely: 1) existing precedent, as the second phenomenon has been directly observed in Pallister-Killian syndrome, where the percentage of abnormal cells decreases with age in blood but not fibroblasts (Conlin et al. 2012), and tissue-limited mosaicism has been observed in mosaic tetrasomies of chromosomes 5p, 8p, 9p and 18p (Choo et al. 2002); 2) the clonality of events observed in both blood and saliva is not greater than the clonality of events in only saliva, which would be expected if events seen across tissue arose earlier in development; 3) both observed LOH events are shared between tissues but only 1 of 9 CNV events are shared between tissues, perhaps suggesting increased pathogenicity of CNV events compared to copy-neutral events, thus more likely to be negatively selected in blood. Given these considerations underlying the disparity in tissue-type, and the observation that the majority of observed abnormalities were detected in saliva but not blood, it is possible that, compared to the sampling of saliva, the sampling of blood could lead to a substantial loss of power, possibly less than 50% power, to detect pathogenic structural mosaicism, resulting in missed diagnoses. Studying the saliva tissue in these children permitted the identification of their mosaic abnormalities and ended for them and their families, their quest for diagnosis.

Additional work is required to investigate for which developmental disorders tissue-limited mosaicism is common. Another intriguing question regarding tissue distribution is the relationship between clonality and pathogenicity. While mosaicism limited to a small number of cells is unlikely to cause developmental disorders, it is conceivable that low-level mosaicism present in a vulnerable tissue, such as white matter neurons, may have clinical consequences. More work is needed to address this question, including more extensive analysis of the tissue distribution of mosaicism, for example, by analysing diverse tissues sampled from all three germ layers, and assays with improved resolution, allowing single or oligo-cell sequencing. The availability of more sensitive detection methods will improve the detection of a larger fraction of events limited to a single tissue.

Next generation sequencing, in the form of exome and genome sequencing, can be harnessed to detect a wide range of mutations, including, as presented here, mosaic structural abnormalities. Given that sequencing costs continue to decline and the multifaceted detection capabilities of exome data, it may be that exome sequencing will supersede microarray technology as a first-line test for developmental disorders. Widespread incorporation of high-depth exome and whole genome sequencing will revolutionise our understanding of the extent of mosaicism in the body and better define the relationship of mosaicism and disease.

## METHODS

### MrMosaic

Implementing mosaic detection requires generating an input file and executing the algorithm; the latter consists of several steps: statistical testing, segmentation, filtering, and results visualisation. ‘BAF’ is used below as an alias for ‘non-reference proportion’. The input data for MrMosaic consist of genomic loci with measured B_dev_ values, C_dev_ values, and genotypes, stored in a tab-delimited file. The loci selected were di-allelic single-nucleotide polymorphic (1%-99% MAFs among European individuals in the UK10K^42^ project) autosomal positions. For exome analysis, only loci overlapping targeted regions of the exome design were used. At these loci, B_dev_ and C_dev_ values were calculated as described in the following two paragraphs.

B_dev_ values were generated using the following method: the identity of the alleles at each locus is extracted using fast_pileup function in the perl module Bio::DB::Sam (Stajich et al. 2002), using high-quality reads (removal criteria: below base quality Q10, below mapping quality Q10, improper pairs, soft-or hard-clipped reads) and BAF was calculated as the number of reference bases divided by the total of reference bases and non-reference bases. Heterozygous sites were defined as loci with a BAF between 0.06 and 0.94, inclusive. The B_dev_ is calculated at heterozygous sites as the absolute difference between the BAF and 0.5. Only loci with sufficient read coverage (at least 7 reads) are used for analysis.

C_dev_ values were generated using the following method: read depths from each target region was collected, the log2 ratio for that target region was calculated by comparing its read depth to a reference read depth, where the reference value was defined as the median read depth among the distribution of read depths at that target region from dozens of highly correlated samples. This log_2_ ratio was normalised based on several covariates pertaining to each target region (covariates included were: GC-content, hybridisation melting temperature, delta free energy (Fitzgerald et al. 2014)). Lastly, using the Aberration Detection Algorithm v2 (ADM2) method by Agilent^®^ a final error-weighted value, is produced, which we use as the C_dev_ value.

The statistical testing step of the MrMosaic algorithm begins by data smoothing, using a rolling median width of 5) across heterozygote and homozygous sites, so as to utilize the depth information in homozygous sites to reduce variance. From this point forward, only heterozygote sites are considered, as mosaic abnormalities do not affect B_dev_ of homozygous loci. Statistical testing assesses whether a given locus is significantly deviated from the B_dev_ and C_dev_ means given the null hypothesis of no chromosomal abnormality. At every heterozygote site we compute two Mann Whitney U tests, one for Bdev and one for Cdev, testing the alternative hypothesis that the distribution of the metric in the neighborhood of the chosen site is greater has a higher median rank) than the distribution of the background. We use 10,000 randomly selected sites, from all autosomes excluding the current chromosome, as the background population. In order to account for non-uniform spacing of the data points we apply a distance-weighted resampling scheme, to down-weight distant points from the chosen site. The tri-cube distance, inspired by Loess smoothing, was chosen as a decay function for the resampling weights and considers data points up to 0.5 Mb upstream and downstream of the given position. An equal number of data points is then sampled around the chosen site and from the background (n=100) and the Mann-Whitney U test is performed. Finally, we combine the p values of the two statistical tests (one for B_dev_ and C_dev_) for every position using Fisher’s Omnibus method.

The segmentation step operates on the combined p value generated above. Segmentation is performed using the GADA algorithm (González et al. 2011), using the parameters values as follows: SBL step: maxit of 1e7; Backward Elimination step: T value of 10 and MinSegLen value of 15. This step generates contiguous segments of putative chromosomal abnormalities. Segments in close proximity (within 1Mb) that show the same signal direction (loss, gain, LOH) are merged to reduce over-segmentation.

The filtering step is required to assess which of the segments generated above are likely reflective of true mosaicism. While testing MrMosaic in exome simulation analyses we observed that true-positive detections (those overlapping simulated events) tended to be larger greater number of probes) and have stronger evidence of deviation GADA amplification value) than putative segments that did not overlap simulated regions i.e. false-positive, spurious calls) (Supplementary Figs. 22-24). We captured these two features in a scoring metric calculated from the cumulative empirical distribution functions for ‘number of probes’ and ‘GADA amplification value’ of false-positive segments, and assessed the composite probability that a given segment comes from these distributions, such that: Mscore = abs(−log_2_(x) + −log_2_(y)) where x and y refer to these empirical cumulative distribution functions. Thus, the Mscore is a quality-control metric derived by combining the size and signal-strength of detections. We used the Mscore to filter those events least likely to represent false positives. We selected events with an Mscore of 8 or greater for analysis because we observed that this appeared to provide a good balance between sensitivity and specificity (Supplementary Figure 24).

The visualisation step generates a detection table and detection plots. The detection table consists of mosaic abnormalities detected and contains the following data: chromosome, start_position, end_position, log2ratio_of_segment, bdev_of_segment, clonality, type, number_of_probes, GADA_amplification, p_val_nprobes, p_val_GADA_amplification, Mscore. Event clonality was calculated by assessing the type of mosaic event based on LRR and converting the bdev value to clonality based on the type of event Supplementary Table^9^). The detection plots are png files showing the loci and BAF and C_dev_ data for each chromosome in which a mosaic abnormality is detected, as well as a genome-wide lattice plot using the data for all chromosomes.

MrMosaic is primarily written in the R language, available as an open source tool at https://github.com/asifrim/mrmosaic. The algorithm can be used in multi-threaded mode to facilitate whole genome analysis. Analysis of a single whole exome using a single thread was completed in 15 minutes when tested using a single core of an Intel Xeon 2.67Ghz processor and 500 Mb of RAM. Whole genome analysis using 24 cores required 30 Gb of RAM and 7 hours. Whole genome analysis can be substantially shortened if the number of sliding windows is reduced or the window size is increased.

### Simulating Mosaicism

We devised a series of simulation experiments to assess MrMosaic performance for various events, across type (LOH, gains, losses), clonalities, sequencing depths, platforms (whole-exome (WE) and whole-genome (WG)) and to compare performance to the MAD method. We compared performance to a modified version of MAD we adapted to enable more flexible execution in a parallel-computing environment, but identical with respect to statistical methods.

The simulation method consisted of these steps: (1) loci selection, (2) calculating depth at these loci, (3) parameter space and number of trials, (4) adjusting read depth in simulated regions, (5) calculating final real depth, (6) selecting sites based on minimum depth, (7) calculating relative copy-number, (8) assigning genotypes, (9) calculating the BAF for each site, (10) calculating performance. Steps 1-3 differed between the WES and WGS simulations and are described first below. The remaining steps 4-10 were executed consistently for WES and WGS simulations and are described next.

For WES simulations, loci selection (1) was based on di-allelic single nucleotide polymorphic positions (between 1% and 99% UK10K (Walter et al. 2015) European minor allele frequency) in the V3 version of the target-region design. To calculate depth at these loci (2), at each locus *i*, baseline sequence read depth (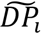 for these sites was defined as the median of the read depth distribution among 100 parental exomes for each site, considering only high-quality reads (mapQ >= 10, baseQ >=10, properly mapped read-pairs), where parental exomes had a mean average sequencing output of 67x (calculated where x was the number of QC-passed & mapped reads without read-duplicates * 75 bp read length / 96 Mb targeted bp). The parameter space (3) consisted of the following: target average sequencing coverage (in x) ∈ {50, 75, 100}, event clonality *m* ∈ {0.25, 0.375, 0.5, 0.75}, type 6 {loss, gain, LOH}, and size ∈ {2e6, 5e6, 1e7, 2e7}. Two hundred trials (4) were conducted per parameter combination for a total of 36,000 simulations.

For WGS simulations, the loci selection (1) was based on di-allelci single nucleotide polymorphic (1% – 99% European MAFs from the 1000 genomes project (Abecasis et al. 2012) May-2013 release) autosomal positions. To calculate depth at these loci (2), we calculated a scaling factor for each locus based on the median read depth of the first two median absolute deviations of the distribution of coverage for that site seen across 2,500 low-coverage samples in the 1000 genomes project (Abecasis et al. 2012). A site-specific scaling factor was calculated as the deviation of each site’s read depth from the average read depth across all polymorphic positions. Simulation depth was defined at each site as the desired simulation coverage multiplied by site-specific scaling factor. The parameter space 3) consisted of two experiments: 1) average genome coverage of 25x, event clonality *m* ∈ {0.25, 0.375, 0.5, 0.75}, type {loss, gain, LOH}, and size (Mb) ∈ {1e5, 2e6, 5e6}; and 2) 5 Mb 50% clonality event captured at average genome coverages (in x) ∈ {30, 40, 50, 60} for the three mosaic types {loss, gain, LOH}. One hundred trials (4) were conducted per WGS simulation.

The remaining simulations steps 4-10 described below were performed consistently for WES and WGS simulations. For each simulation a single mosaic event was introduced into each simulation trial. The adjustment of read depth in simulated regions 4) was performed using a scaling factor based on the type and clonality of the simulated event, *m*, while sites not overlapping copy-number simulated events would not undergo this scaling step (Supplementary Table 4). To calculate the final simulated read depth (5) for each site *i*(*SDP_i_*), we sampled from a Poisson distribution with *λ_i_* equal to the scaled read depth. Only positions with a final read depth (6) of at least 7 were included for analysis. Relative copy-number (7) was defined as log_2_ of the ratio of the final read depth to the baseline read depth.

The assignment of genotypes (8) (AA, AB, or BB) at each position *i* was randomly determined based on the site’s minor allele frequency, which was used in a multinomial function with probabilities corresponding to Hardy Weinberg-assumed genotype proportions (p^2^, 2pq, q^2^). To calculating the BAF for each heterozygote at site *i* (9), we adjusted the expected heterozygote proportion of 0.5 with respect to the chosen event type and clonality, and sampling from a binomial distribution given this adjusted proportion and the simulated read depth at *i.* BAFs for homozygote reference (AA) and non-reference (BB) sites were chosen by sampling from a binomial distribution with p=0.01 or p=0.99 respectively and the simulated read depth at *i*.

MrMosaic and MAD were applied on the simulated WES and WGS samples generated by the above procedure and performance was measured using precision-recall metrics (10). A ‘success’ in a trial was considered a detection overlapping the simulated mosaic event. Precision was calculated as the number of successes divided by the number of detections. Recall was defined as the proportion of trials with a success.

### Description of Samples & Sequencing

The samples used in this analysis derived from the Deciphering Developmental Disorders study, a proband two-parent trio-based investigation of children with undiagnosed developmental disorders from the UK and Ireland (King et al. 2015; Firth and Wright 2011; Wright et al. 2014; Fitzgerald et al. 2014). DNA was extracted from blood and saliva and was processed at the Wellcome Trust Sanger Institute by array CGH and exome sequencing. There were 4,926 DNA samples analysed in this study from 4,911 children, as some children were analysed using both blood and saliva. The majority, 3,260 of 4,926 (66%) of the DNA samples were extracted from saliva.

DNA was enriched using a Agilent^®^ exome kit, based on the Agilent Sanger Exome V3 or V5 backbone and augmented with 5 Mb of additional custom content (Agilent Human All Exon V3+/V5+, ELID # C0338371). An ‘extended target region’ workspace was defined by padding the 5′ and 3′ termini of each target region by 100-bp yielding a total analyzed genome size of approximately 90 Mb. Sequencing was performed using the Illumina^®^ HiSeq 2500 platform with a target of at least 50× mean coverage using paired-end sequence reads of 75-bp read-length. Measured exome coverage ranged from 14× to 155× with a mean of 69× (Supplementary Figure 24). Alignment to the reference genome GRCh37-hs37d was performed by bwa version 0.5.9 (Li and Durbin 2009) and saved in BAM-format files (Li et al. 2009).

Additionally, two exome samples were processed *post hoc* from saliva after SNP genotyping chip analysis showed mosaicism was present in saliva but absent in blood. These two exome samples and the exome sample with suspected revertant mosaicism were processed separately from the exome experiment described in the previous paragraph. For these three exomes, the Agilent Sanger Exome V5 target kit was used, and sequence depth ranged from 387x – 455x coverage (reads = {465,522,627, 483,098,826, 549,766,632} * 75bp read-length/90e6 target-region-size). The sample with suspected underlying mosaic reversion had 549,224,891 QC-passed & mapped reads, and 57,165,328 duplicates, and therefore had a mapped read coverage of 410x ((549,224,891-57,165,328))(* 75/90e6).

For the sample for which whole genome sequencing data were generated, sequencing was performed using an Illumina^®^ X-Ten sequencing machine. Library fragments of 450-bp insert-size were used and paired-end 151-bp read-length sequence reads were generated. Alignment to the reference genome GRCh37-hs37d was performed by bwa mem^47^ version 0.7.12. Average coverage was calculated using samtools flagstat as the number of QC-passed mapped-reads without duplicates using 151 bp read-lengths in a 3Gb genome: (616,151,282 −124,325,581) *151/3e9 = 24.8×. Rearrangement analysis was carried out using Breakdancer v1.0 Chen et al. 2009).

### Additional filtering implemented in addition to Mscore quality score

Some events with very high Mscores appeared to represent real, but constitutive, abnormalities. There were two failure modes we identified: constitutive duplications and homozygosity by descent (HBD). Constitutive duplications genuinely produce strong signals in MrMosaic, but also constitutive deletion and ROH events may produce putative detections if individual probes had mapping artefacts that resulted in spurious signals. We used bcftools roh (manuscript in preparation) to identify and filter HBD regions and flagged as suspicious events with greater than 25% reciprocal overlap with CNVs detected through constitutive copy-number detection. In addition, we observed several recurrent putative detections, especially prevalent in pericentromeric and acrocentric regions that appeared spurious on the basis of inconsistencies between BAF and LRR, and we filtered such systematic errors by filtering putative mosaic events seen in more than 2.5% of samples.

### SNP genotyping chip validation

Illumina^®^ HumanOmniExpress-24 Beadchips (713,014 markers) were used. Illumina GenomeStudio software was used to generate log R ratio and BAF metrics and Illumina^®^ Gencall software was used to calculate genotypes. Structural mosaic detection was performed using MAD (González et al. 2011). Initial mosaic events were merged if events were within 1 Mb, and were the same type (loss, gain, or LOH) of mosaic event. Results were plotted using custom R code.

## DATA ACCESS

MrMosaic source code can be found here: https://github.com/asifrim/mrmosaic.

The complete raw exome sequencing data is publicly available on the European Genome-phenome Archive (EGA) after Data Access Committee (DAC) approval. The study accession ID is: EGAS00001000775.

## ACKNOWLEDGMENTS

This study relied on the generous participation of DDD patients and their parents. We are grateful to the DDD informatics and HGI pipeline staff for generating the data, DDD laboratory staff for sample handling, and Sanger genotyping core for running the validation arrays. Yanick Crow, Helen Firth, David Fitzpatrick, and Wendy Jones provided invaluable clinical expertise. We thank Jeff Barrett for his sharp, constructive feedback. Rolph Pfundt and James Lupski aided the interpretation of the revertant mosaic mutation. The DDD is supported by the Health Innovation Challenge Fund (HICF-1009-003), a parallel funding partnership between the Wellcome Trust and the Department of Health, and the Wellcome Trust Sanger Institute (WT098051).

## AUTHOR CONTRIBUTIONS

DK, AS and MH developed the algorithm and wrote the manuscript; TF generated the ADM2 exome scores; YC, EH, TH, SM, SM, MS, ST, PV recruited and phenotyped the DDD patients with detected mosaicism.

